# Phylogenetic diversity of the carbon monoxide-utilizing prokaryotes and their divergent carbon monoxide metabolisms in the human gut microbiome

**DOI:** 10.1101/2023.10.23.563559

**Authors:** Yuka Adachi Katayama, Ryoma Kamikawa, Takashi Yoshida

**Affiliations:** Graduate School of Agriculture, Kyoto University, Kitashirakawa Oiwake-cho, Sakyo-ku, Kyoto 606-8502, Japan

**Keywords:** Human gut microbiome, Carbon monoxide-utilizing prokaryotes, Wood-Ljungdahl pathway, Nickel-containing carbon monoxide dehydrogenase, *Blautia*, *Veillonella*

## Abstract

Although the production of toxic CO within the human body has been detected, only a few CO-utilizing prokaryotes (CO utilizers) have been reported in the human gut, and their phylogenetic and physiological diversity remains unclear. Here, we unveiled more than thousand representative genomes originating from previously unexplored potential CO-utilizing prokaryotes, which contain CO dehydrogenase (CODH) genes. More than half of CODH-bearing prokaryotes possess genes for the autotrophic Wood–Ljungdahl pathway (WLP). However, 79% of these prokaryotes commonly lack a key gene for WLP, which encodes enzyme that synthesizes formate from CO_2_ and reductants such as H_2_, suggesting that they share a degenerated WLP. Instead, many were predicted to possess an alternative way of synthesizing formate from pyruvate, which is a product of glycolysis. In addition to degenerated WLP, seven genes neighboring the CODH gene were found, which may reflect diverse utilization of CO in the human gut. Our findings reveal the unique and diverse nature of CO metabolism in the human gut microbiome, suggesting its potential contribution to CO consumption and gut homeostasis.

**Impact statement:** Carbon monoxide (CO)-utilizing prokaryotes mitigate the toxic impact of CO by consuming it as energy and/or carbon sources. In addition to various environments, CO is also produced via multiple routes, such as heme degradation, in the human body and accumulates in the gut. Revealing CO-utilizing prokaryotes and their CO metabolisms in the human gut would contribute to gaining insight into how microbial community functions are involved in maintaining human gut homeostasis. Nevertheless, the limited number of CO utilizers in the human gut microbiome have been reported. In our study, a significant proportion of human gut microbial genomes belonging to diverse phyla were revealed to be of potential CO-utilizing prokaryotes. Additionally, the majority of CO-utilizing prokaryote genomes in the human gut have potentially remodeled the Wood– Ljungdahl pathway (WLP), one of the most well-known autotrophic pathways, to the degenerated, heterotrophic form. Moreover, there were seven other genes neighboring CODH in the human gut CO-utilizers, suggesting various CO utilization. Our findings would pave the way for future explorations into microbial metabolic adaptations and their implications for human health.

**Data summary:** The human gut prokaryote genomes were downloaded from HumGut database (Hiseni et al. 2021; https://arken.nmbu.no/~larssn/humgut/). The accession numbers of CODH/ACS-bearing genomes from environments without host-association (Inoue *et al*., 2022) are listed in Table S1. Metatranscriptomic datasets were downloaded from the NCBI Sequence Read Archive (SRA) under the Bioproject accession numbers PRJNA354235 and PRJNA707065 and their accession IDs are listed in Table S2.

## Introduction

Carbon monoxide (CO) is a colorless, odorless, and tasteless gas that is toxic to many organisms (Ernst and Zibrak 1998). CO has a 200–250 times higher affinity for hemoproteins, such as hemoglobin and cytochrome *c*, than oxygen. Therefore, exposure to >100 ppm of CO inhibits oxygen transport and respiration, resulting in severe effects on human health (Ernst and Zibrak 1998; Alonso *et al*. 2003; Prockop and Chichkova 2007). Nevertheless, the human body produces 16.4–160 μmol, which correspond to 0.5–4.5 ppm, of CO per hour on average, primarily through heme degradation using heme oxygenase-1 (Morse and Choi 2005; Onyiah *et al*. 2014; Hopper *et al*. 2020). From the human body and exogenous sources, CO accumulates in the gastrointestinal tract, potentially influencing the composition of intestinal microbiota (Hopper et al. 2020).

Prokaryotes utilizing CO are called CO utilizers and have been found in various environments, including soil (Suzuki *et al*. 1983), lake (Leigh *et al*. 1981), freshwater lake sediments (Imaura *et al*. 2023), bioreactors (Hattori *et al*. 2000), hot springs (Svetlichny *et al*. 1991), deep hydrothermal vents (Moussard *et al*. 2004), and even gut environment (Trischler *et al*. 2022). All CO utilizers possess CO dehydrogenase (CODH), which catalyzes the reversible conversion of CO_2_ and CO (CO_2_ +[2H^+^ +[2e^−^ <=> CO[+[H_2_O) (Oelgeschläger and Rother 2008; Sokolova *et al*. 2009). CODHs are physiologically and structurally diverse enzymes that form two major groups, namely nickel- and molybdenum-containing CODHs (Ni-CODH and Mo-CODH); of which, Ni-CODH works under anaerobic conditions (Ragsdale 2004). The catalytic subunit of Mo-CODH (CoxL) is phylogenetically classified into two groups, form I and II, whereas that of Ni-CODH (CooS and CdhA) is phylogenetically and structurally classified into eight distinct clades, A to H, and mini-CODH (Inoue *et al*. 2019, 2022).

Through reactions catalyzed by the abovementioned CODHs, CO utilizers utilize CO as a carbon source and/or as the low redox potential (E^0^′=[−520 mV) for carbon fixation and/or energy conservation in various CO metabolizing pathways and respiratory chains. For instance, through the reduction and oxidation of the ferredoxin-like protein CooF and flavin adenine dinucleotide-dependent NAD(P) oxidoreductase (FNOR), the reducing power of CO is supplied to NAD^+^, thereby generating NADH, which is available for various respiration and biosynthesis reactions (Tian *et al*. 2016; Inoue *et al*. 2019; Slobodkin *et al*. 2019). The reducing power of CO is also supplied to various terminal electron acceptors, including proton, nitrate, and oxygen, in the electron transport chains to generate an electrochemical gradient for ATP synthesis (Oelgeschläger and Rother 2008; Slobodkin *et al*. 2019; Adachi *et al*. 2020).

Among the CO-mediated CO_2_ fixation pathways, the Wood-Lijungdahl pathway (WLP; reductive acetyl-CoA pathway) has been best studied. The WLP is recognized as the most ancient autotrophic pathway, which was possessed by the last universal common ancestor (LUCA) (Schuchmann and Müller 2014; Weiss *et al*. 2016; Adam *et al*. 2018). This pathway consists of two branches, carbonyl and methyl branches (Ragsdale and Pierce 2008; Adam *et al*. 2018). In the carbonyl branch, CO_2_ is reduced to CO by Ni-CODH and incorporated into the carbonyl moiety of acetyl-CoA. Alternatively, CO can be obtained directly from the surrounding environment and incorporated into the carbonyl branch (Ragsdale 2008). In the methyl branch, formate is synthesized from CO_2_ and reductants such as H_2_ by formate dehydrogenase (Fdh), which are then reduced stepwise to a methyl moiety (CH_3_-) by formyl-tetrahydrofolate synthase (Fhs), methylene-tetrahydrofolate dehydrogenase/cyclohydrolase (FolD), and methylene-tetrahydrofolate reductase (MetF). The generated carbonyl and methyl moieties are incorporated into acetyl-CoA along with coenzyme A (CoA) by acetyl-CoA synthase (ACS), which forms a complex with Ni-CODH (CODH/ACS). Acetyl-CoA is then converted into various compounds such as short-chain fatty acids (SCFA), including acetate, by CO utilizers (Miller and Wolin 1996; Trischler *et al*. 2022). In many of these acetogenic CO utilizers, WLP is linked to energy conservation using Rnf complexes and/or energy-converting hydrogenase (Ech), where the oxidation of reduced ferredoxin is coupled with the reduction of NAD^+^ and protons, respectively (Diender *et al*. 2015; Schoelmerich and Müller 2019).

Although the CO-utilizing abilities of human gut prokaryotes have not yet been fully elucidated, accumulating evidence suggests that CO utilizers are present in the human gut microbiome. Recently, two human gut bacteria, *Blautia luti* and *B. wexlerae*, were found to consume CO via the WLP when formate was added to the medium (Trischler *et al*. 2022). Furthermore, several gastrointestinal prokaryotes such as *Clostridioides* and *Marvinbryantia* possess genes for WLP (Wolin *et al*. 2003; Yao *et al*. 2023). However, the diversity of CO utilizers and their metabolic pathways in the human gut remain poorly understood.

With the advancement of sequencing technologies and bioinformatic tools, extensive databases of human gut prokaryotic genomes have been established. Almeida *et al*. (2021) created a Unified Human Gastrointestinal Genome (UHGG) collection comprising 204,938 non-redundant genomes. Hiseni *et al*. (2021) screened more than 5,700 healthy human gut metagenomes and constructed HumGut, a genome collection containing over 381,000 genomes. Analyzing these databases would deepen our understanding of human intestinal prokaryotes, including CO utilizers and uncultured prokaryotes, which are dominant in the human gut (Almeida *et al*. 2019, 2021; Omae *et al*. 2019). In this study, we used the HumGut database to investigate CO utilizers and their CO-utilizing pathways in the human gut microbiome.

## Methods

### Genome sources

Publicly available genome collection, HumGut (Hiseni *et al*. 2021), was used as a source of human gut prokaryote genomes (downloaded from: https://arken.nmbu.no/~larssn/humgut/). In the database, >381,000 genomes detected from 3,534 healthy human gut metagenome samples were clustered into 30,691 representative genomes at 97.5% sequence identity. The previously reported WLP-bearing bacterial genomes without host asscociation (Inoue *et al*. 2022) were utilized and listed in Table S1. Metatranscriptomic datasets derived from 110 different healthy individuals were downloaded from the NCBI Sequence Read Archive (SRA) under the Bioproject accession numbers PRJNA354235 and PRJNA707065 (Table S2).

### Detection of *cooS* and *coxL* in human gut prokaryote genomes

CODH-bearing genomes were searched from 30,691 representative human gut prokaryotic genomes, using *cooS* and *coxL* as genetic markers for Ni- and Mo-CODHs, respectively. The homology search was performed using NCBI blast+ 2.13.0 with cutoffs of an E-value of 10^−10^, a sequence length of 200 aa, and a sequence identity of 30%, using the following amino acid sequences as queries: clades A–H CODHs and mini CODH from *Methanosarcina barkeri* CODH/ACS α subunit (WP_011305243.1), *Butyrivibrio* sp. CooS (WP_026514536.1), *Clostridium novyi* CooS (WP_039226206.1), *Carboxydothermus hydrogenoformans* CooSV (WP_011342982.1), *Thermococcus onnurineus* CooS (WP_012571978.1), *C. hydrogenoformans* CooSII (WP_011343033.1), *Deltaproteobacteria bacterium* RBG_16_49_23 CooS (OGP75751.1), Candidatus Atribacteria bacterium HGW-Atribacteria-1 CooS (PKP59679.1), and *Thermosinus carboxydivorans* mini CooS (WP_007288589.1), respectively (Inoue *et al*., 2019, 2022; Techtmann *et al*., 2012). The Ni-CODH clades of the detected CooS were determined by their amino acid sequence identity with representative CooS sequences. The detected CooS/CoxL sequences were aligned using MAFFT v7.487 with the E-INS-i application (Katoh and Standley 2013). To examine the motifs of CooS, previously used criteria (Inoue *et al*. 2019) were adopted: no amino acid substitutions in two C-clusters comprising Ni, Fe, and S; a B-cluster comprising cubane-type 4Fe-4S; and a D-cluster comprising an additional 4Fe-4S at the subunit interface (Dobbek et al., 2001; Doukov *et al*. 2002; Inoue *et al*. 2019). A similar analysis was performed for CoxL with slight modifications. A homology search was performed using *Oligotropha carboxidovorans* CoxL (WP_013913730.1) as the query. The sequences that conserved the Form I CoxL active site (AYRCSFR) (King 2003) alone were selected. The detected Ni-CODHs are listed in Table S3. All genomes containing Ni-CODHs are listed in Table S4.

### Phylogenetic tree construction

The taxonomic assignment of 30,691 prokaryotic genomes was performed using GTDB-tk v2.1.1, with the reference data version R207 (Chaumeil *et al*. 2019). The names of the phyla were corrected according to the method described by Oren and Garrity (2021). Genome distances among the *cooS*-containing genomes were calculated using Mashtree v1.2.0 with 1,000 replicates and a minimum depth value of 0 (Ondov *et al*. 2016; Katz *et al*. 2019). The obtained tree was then visualized using the R package, ggtree v3.2.1 (Yu *et al*. 2017). Of the 3,534 human gut metagenome samples (Hiseni *et al*. 2021), the number of metagenome samples, from which the bacterial genomes were detected with 95% or higher identity, was displayed as a heatmap on the outer layer of the phylogenetic tree.

### Genome-based prediction of metabolic functions

To characterize the metabolic functions of the CO utilizer in the human gut, three groups of genomes were analyzed using METABOLIC v4.0 (Zhou *et al*. 2022). The first group comprised the CODH-bearing genomes detected in the HumGut database. The second group comprised the non-CODH-bearing genomes of human gut prokaryotes belonging to a family or order with CODH-bearing genomes, that were detected in the HumGut database. The third group comprised the CODH/ACS-bearing bacteria found in environments other than the intestine. The proportions of genomes encoding functions in each genus were calculated for each group. From the HumGut database, we selected Ni-CODH-lacking bacterial genomes belonging to orders or families containing Ni-CODH-bearing bacteria, resulting in 194 genomes (the second group). We also utilized Ni-CODH-bearing bacterial MAG datasets constructed by Inoue et al. (2022). Among the genomes, only the 554 genomes with bacterial CODH/ACS and without “host-associated” tag were used in this analysis. The accession ID of the genomes are listed in Table S1.

### Characterization of gene compositions in the *cooS* genomic contexts

Protein-coding genes in the genomes were predicted using Prodigal v2.6.3, with default settings (Hyatt *et al*. 2012). The gene functions were then annotated to the protein coding sequences using eggNOG-mapper v2.1.5 (Cantalapiedra *et al*. 2021). For double-checking, a homology search of the protein-coding sequences against the clusters of orthologous gene (COG) database was performed using DIAMOND BLASTp v 2.0.12 (Buchfink *et al*. 2021). The KEGG orthology (KO) and COGs of 15 genes located in the upstream and downstream of the CooS loci were then extracted and are listed in Tables S5 and S6, respectively. Previously reported CODH-related COGs were also referred: AcsB (COG1614), CooF (COG0437 or COG1142), FNOR (COG1251), ECH (COG3260 and COG3261), and ABC transporter (COG0600, COG1116, and COG0715) (Inoue *et al*. 2019). The gene maps around *cooS* were visualized using the R package gggenomes v0.9.5.9000 (Hackl and Ankenbrand 2022).

### Genomic-based predictions of the degenerated WLP

In the present study, the gene composition of the CODH/ACS-bearing genomes was analyzed. Genomes that possessed both CooS and AcsB (K14138) were termed CODH/ACS-bearing genomes. All the KOs annotated to bacterial genomes in the human gut and other environments were listed, and the presence of each KO was checked in each genome to calculate the proportion of genomes carrying a given gene in each genome group. When two or more KOs were annotated to a single gene, all annotated KOs were considered for calculating the proportion. Since K00656 (PflD) contains members of PFL-like proteins without PFL activity, proteins annotated as K00656 were aligned using MAFFT v7.487 with the E-INS-i application, and only the sequences that conserved the Cys-Cys active sites (Sawers and Watson 1998) were regarded as potential PFL-coding sequences. The proportions of the genomes carrying the given genes are listed in Table S7. Especially, the proportion of genomes carrying the following genes were extracted from the obtained results: CO metabolism-related genes (*cooC*, K07321; *cooF*, K00196), ACS subunits (*acsB–E*, K14138, K00197, K00194, K15023; *cdhC*, K00193), WLP (*fhs*, K01938; *folD*, K01491; *metF*, K00297; *metV*, K25007), Rnf subunits (*rnfA–G*, KO03612–17; *rnfC2*, K25008), *ech* (K15830, K15832), Hdr (*hdrA–C*, K03388–90), Mvh (*mvhD*, K14127), and PFOR (*por*, K03737; *porA–D*, K00169–72). In addition, formate-related genes were searched in the KO database and KEGG reaction database by inserting the keyword, “formate.” Hydrogenases were omitted from the results and other genes were checked as the formate-related proteins: formate transporters (FocB, K03459; FocA, K06212; OxlT, K08177; YfdC, K21990; FdhC, and K21993), formate dehydrogenase catalytic subunits (FdhA, K05299; FdhF, K22015; and FdoG, K00123) and the other subunits of formate dehydrogenase (K00122–27, K22515, and K22516), PFL (K04069 and K00656), and other 55 genes. Genes that were not present in any genome were removed from the figure.

### Taxonomic and functional profiling of metatranscriptomes

Publicly available metatranscriptome datasets of healthy human were retrieved from the SRA database. Among the 110 datasets, 96 datasets were sourced from BioProject PRJNA354235, which includes samples from healthy adult men from United States aged 40–75 years old (Abu-Ali *et al*. 2018), while 14 datasets were sourced from BioProject PRJNA707065, which includes samples from mothers and their six-month-old infants from New Zealand or United Kingdom under the NiPPeR Study (https://www.nipperstudy.com/). Sequence reads were processed using KneadData v0.12.0 (http://huttenhower.sph.harvard.edu/kneaddata). In KneadData, the quality control pipeline was set as follows: FastQC v0.12.1 (https://www.bioinformatics.babraham.ac.uk/projects/fastqc/) was employed before and after quality controls, adaptor trimming was performed with default settings, reads were filtered by Trimmomatic v0.35 (Bolger, Lohse and Usadel 2014) with a sliding window sizes of four and a minimum quality score of 20, Tandem Repeats Finder (TRF) v4.09 (Benson 1999) was employed with default parameters, and human host RNA (Hg38) and rRNA sequences (Silva v128) were removed using Bowtie2 v2.5.2 (Langmead and Salzberg 2012). Trimmed non-human reads were subsequently mapped to (i) the 1,380 of human gut non-redundant CODH mRNA sequences identified in this study, (ii) the 30,691 human gut prokaryotic genomes in the HumGut database, and (iii) Fdh genes of *B. schinkii*(JANSWJ010000003.1:269667-271859) and *B. hydrogenotrophica* (NZ_CYXL01000001.1:c262854-260023) using CoverM v0.6.1 (https://github.com/wwood/CoverM).

## Results

### CODH was possessed by 4.2% of human gut prokaryotic genomes

In the present study, we analyzed human gut prokaryotic genomes from the HumGut database (Hiseni *et al*., 2021). The genomes in the HumGut database were clustered into 30,691 genomes with 97.5% sequence identity (Hiseni *et al*., 2021). Prior to exploring CO utilizers in the human gut, 30,691 representative genomes were taxonomically classified using GTDB-tk (Chaumeil *et al*. 2019), and 98% of the gut genomes belonged to the phyla Bacillota (19,877 genomes), Bacteroidota (4,011), Actinobacteriota (3,676), Pseudomonadota (1,905), Campylobacterota (227), and Thermodesulfobacteriota (259) (Fig. 1a). The dominant orders in the Bacillota phylum were Oscillospirales (8,642), Lachnospirales (3,608), Christensenellales (1,476), Lactobacillales (1,423), and Veillonellales (1,279) (Fig. 1a).

**Fig. 1.**
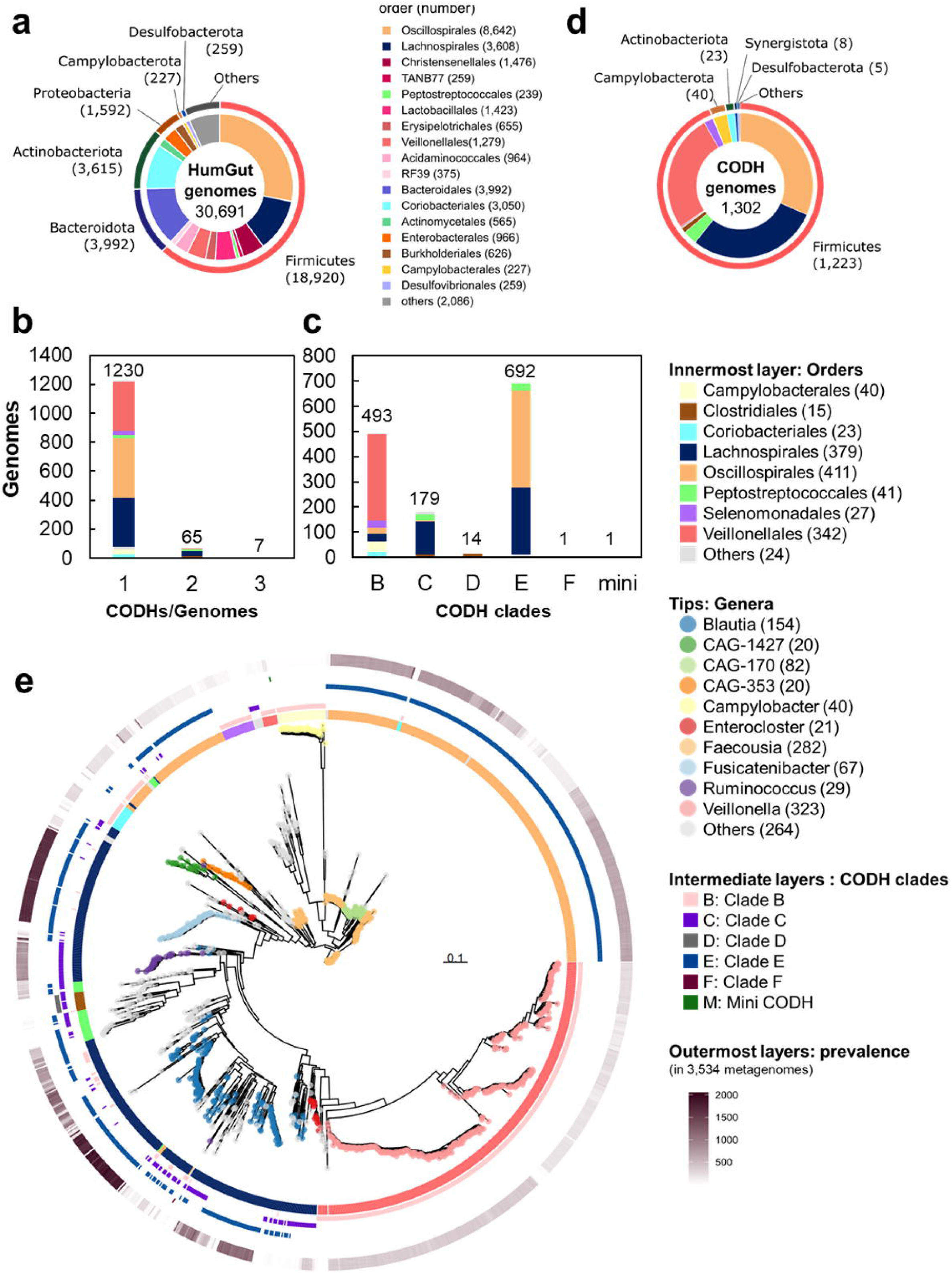
The CODH-bearing prokaryotic genomes in the human gut microbiome. (a) Taxonomic classification of publicly available 30,691 human gut prokaryotic genomes. The genomes were the representatives of more than 0.38 million genomes, clustered with 97.5% nucleotide sequence identity (Hiseni *et al*., 2021). The numbers in parentheses show those of genomes. (b) The number of genomes that possess 1–3 CODHs. No genome possessed four or more CODHs in the used dataset. (c) The number of genomes that possess a CODH belonging to any of the six clades. (d) Taxonomic classification of the CODH-bearing genomes. (e) Phylogenetic tree of the CODH-bearing genomes. The tip color represents the genus level classifications. The heat maps around the phylogenetic tree exhibits order level taxonomic classifications, CODH clades B, C, D, E, and F and mini CODH from the inner layer, respectively. The outermost layer shows the prevalence of each genome reported in the previous study (Hiseni *et al*. 2021). The number indicates the metagenome samples where each genome was analyzed in 3,534 metagenome datasets.

To identify CO utilizers in the human gut, genes for the catalytic subunit of Ni-CODH clades A–G (*cdhA* and *cooS*) and mini CODH were used as markers for CO utilizers. As a result of homology search, 2,505 amino acid sequences encoded by 2,202 genomes were detected as candidate Ni-CODHs. Of these, Ni-CODHs were further filtered using the following criterion: no amino acid substitutions or deletions in the conserved motifs of the CODH of the five metal clusters and two acid-base catalysts (Inoue *et al*. 2019). Consequently, 1,380 amino acid sequences in 1,302 genomes were confirmed to conserve Ni-CODH motifs. Most of the genomes contained one Ni-CODH gene (1,231/1,302 CODH-bearing genomes), whereas other genomes contained two or more genes (Fig. 1b). The detected 1,380 Ni-CODHs belonged to clades B (493), C (179), D (14), E (692) and F (1) and mini CODH (1) (Fig. 1c). Although a similar survey was conducted for Mo-CODH using CoxL as a marker, Mo-CODH, which possesses a conserved active site for form I CoxL, was not found in any human gut genome.

Among the 1,302 CODH-bearing genomes, the phylum Bacillota comprised 1, 223 (94%). The remaining 79 genomes were identified as Actinobacteriota, Bacteroidota, Campylobacterota, Thermodesulfobacteriota, Pseudomonadota, Synergistota, and Verrucomicrobiota (Fig. 1d, e). The 1,223 Bacillota- and 79 other phyla-derived genomes belonged to 248 species, 82 genera, and 8 phyla.

Among the 1,223 Bacillota-derived genomes, 379, 341, and 411 belonged to Lachnospirales, Veillonel l ales, and Oscillospirales, respectively; thus, a large proportion of the CODH-bearing genomes were occupied by these three orders (Fig. 1d). CODH-bearing genomes accounted for 10.5%, 26.7%, and 4.8% of the total human gut microbial genomes derived from the orders Lachnospirales, Veillonelalles, and Oscillospirales, respectively. Within the order Lachnospirales, 38 genera possessed CODH, including *Blautia* (154/291; 154 Ni-CODH-bearing genomes within 291 of the total genomes of the genus in the human gut microbiome), *Fusicatenibacter* (67/215), *Ruminococcus* (29/1,194), *Enterocloster* (21/56), *Anaerobutyricum* (12/30), and UMGS1375 (10/11). Of these, *Blautia* was previously identified as possessing functional WLP (Trischler *et al*. 2022) and was the only taxon to possess three CODHs (HumGut ID: 13241, 3275, 5110, 12728, 20698, 11367, and 8982) (Fig. 1b). In *Blautia*, 24 of 154 CODH-bearing genomes contained two or more CODH genes. The Veillonelalles with Ni-CODH genes consisted of three genera: *Veillonella* (323/443), *Megasphaera* (12/96), and F0422 (7/12). *Veillonella* is a ubiquitous bacterium found in the human body, such as in the oral cavity and gut (Rogosa 1964). The order Oscillospirales contains 13 genera that encode Ni-CODHs in their genomes, including *Faecousia* (282/446), CAG-170 (82/179), and CAG-353 (20/60) (Table S4). Although Oscillospirales occupies a higher proportion of the human gut CODH-bearing genome, most of these bacteria are uncultured; therefore, their physiological characteristics are largely unknown. CODH-bearing genomes were also identified in the Bacillota orders Peptostreptococcales (41/239) and Selenomonadales (27/135) as well as in the Actinobacteriota order Coriobacteriales (23/3,050) (Table S4).

To assess the prevalence of potential CO utilizers in humans, we checked for the presence of CODH-bearing genomes among the available 3,534 datasets of healthy human gut metagenomes (Hiseni *et al*., 2021) (Fig. 1e). The CODH-bearing genomes derived from the Lachnospirales genera *Fusicatenibacter* and *Blautia* were present in 48% and 26% of the metagenome datasets, respectively. The Veillonelalles genera *Veillonella* was detected in 8.3% of the metagenome datasets (Fig. 1e, Table S4). The Oscillospirales genera *Faecousia* and CAG-170 were detected in 13% and 18% of the metagenome datasets, respectively. The above CODH-bearing genomes were appeared more prevalent than or as prevalent as the two prominent CODH-lacking Bacteroidales genera, *Phocaeicola* and *Bacteroides* (Hiseni *et al*. 2021), which were detected in the 18% and 31% metagenome datasets on average, respectively. In contrast, some CODH-bearing genomes were present in only a limited number of the human gut metagenome datasets (Fig. 1e, Table S4). For instance, the CODH-bearing genomes from the genus *Eubacterium* in the order Eubacteriales were found in only 0.2% of metagenomes on average.

### Majority of CODH-bearing bacteria were acetate-producing prokaryotes in the human gut

As most of the potential CO utilizers detected in this study have been uncultured and thereby uncharacterized, we reconstructed the genome-based metabolic functions of the potential CO utilizers bearing Ni-CODH genes in the human gut microbiome using METABOLIC v4.0 (Zhou *et al*. 2022). Twenty-three metabolic functions were detected in the potential human gut CO utilizers, and the acetate production function was conserved among them (Fig. S1). In particular, the analysis revealed that 1,162/1,302 Ni-CODH-bearing human gut genomes, which correspond to 73/82 genera, contained genes for acetate production (*acdA*, *ack*, and *pta*) for fermentation, suggesting that most potential CO utilizers produce acetate in the human gut (Fig. S1). Exceptions were the Selenomonadales genera (*Centipeda*, *Mitsuokella*, and *Selenomonas*), whose all 27 genomes lacked the abovementioned genes for acetate production.

To evaluate whether these functions were specific to potential CO utilizers, we compared the metabolic functions of 1,302 Ni-CODH-bearing human gut microbial genomes to those of 194 Ni-CODH-lacking genomes that were closely related to the Ni-CODH-bearing ones. All functions other than CO metabolism were observed in both types of genomes; thus, we could not find any specific metabolic functions in the potential CO utilizers (Fig. S1). This suggests that CO metabolism may be engaged in accessory functions that support the existing functions conserved in both Ni-CODH-bearing and Ni-CODH-lacking human gut microbiomes.

### Eight *cooS* genomic contexts were identified in the human gut prokaryote genomes

Genes that encode proteins involved in CO metabolism are often located close to *cooS* in the genome, enabling the prediction of their physiological roles based on the genomic context (Inoue *et al*. 2019; Matson *et al*. 2011; Techtmann *et al*. 2012). To gain insight into the physiological roles of CO metabolism in the human gut microbiome, we analyzed 15 genes located in the upstream and downstream regions of *cooS* in the 1,302 human gut microbial genomes (Fig. 2, Table S5, S6). Totally, 1,150 KOs and 1,285 COGs were annotated for genes within the 1,380 *cooS* contexts in the 1,302 genomes (Tables S5 and S6). Genomic contexts were manually classified into eight types: WLP, PEPCK, FNOR, ABC transporter, Fe-only hydrogenase, MFS transporter, uncharacterized dehydrogenase, and Cysteine synthase. Below, we describe the estimated physiological functions and detailed phylogenetic distributions of genomic context types (Fig. 2).

**Fig. 2.**
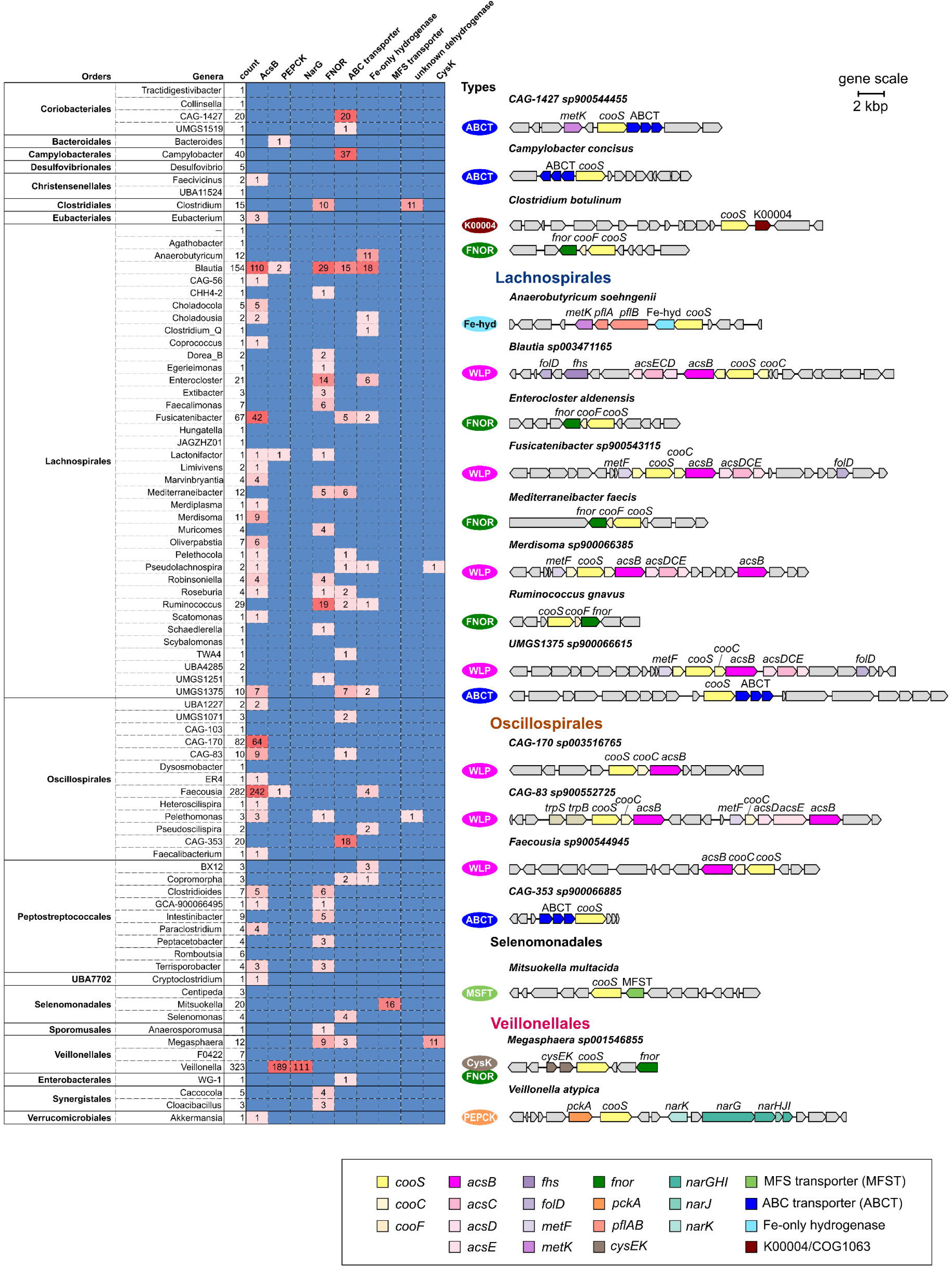
The genomic context of CODH (*cooS*) in the CODH-bearing genomes in the human gut microbiome. The heatmap on the left shows the number of genes detected within the 15 genes upstream and downstream of *cooS* in the genomes of each genus. Genera with more than nine CODH-bearing genomes were selected (red font), and the gene map around *cooS* of the genera is shown on the right. Each box represents protein-coding genes and their coded strands. The gray box represents the others and the hypothetical protein. *cooS*, carbon monoxide dehydrogenase catalytic subunit gene; *cooC*, carbon monoxide dehydrogenase maturation gene; *cooF*, ferredoxin-like protein-coding gene; *acs*, acetyl-CoA synthase gene; *fhs*, formyltetrahydrofolate synthase gene; *folD*, methylene-tetrahydrofolate dehydrogenase/cyclohydrolase gene; *metF*, methylene tetrahydrofolate reductase gene; *metK*, S-adenosylmethionine synthetase gene; *fnor*, flavin adenine dinucleotide-dependent NAD(P) oxidoreductase gene; *pckA*, phosphoenolpyruvate carboxykinase gene. *pfl*, pyruvate: formate lyase gene; *cys*, cysteine synthase gene; *nar*, nitrate reductase gene; MFS transporter, major facilitator superfamily transporter; ABC transporter, ATP-binding cassette transporters.

The WLP type, characterized by the presence of the gene for ACS β subunit (AcsB; K14138, COG1614), was the most prevalent in the human gut microbial genomes, being detected from 593 of the 1,380 *cooS* contexts. The WLP context was observed in Oscillospirales (8/13 genera, that is, eight CODH-bearing genera within the total genera of the order; 323/411 CODH-bearing genomes, that is, 323 CODH genomes within the total genomes of the order), Lachnospirales (18/38 genera, 198/379 genomes), Peptostreptococcales (4/9 genera, 13/41 genomes), and Eubacterales *Eubacterium* (3/3 genomes). All genera conserved the consecutive gene arrangement of *cooS*-*cooC*-*acsB*, except for the Peptostreptococcales genera, which had the gene arrangement of *cooS*-*cooC*-*fhs* and *acsB* at 10 genes downstream of *cooS* (Tables S5 and S6). *acsCDE* was found in the *cooS* contexts of the Lachnospirales and Peptostreptococcus genera but not in seven out of eight Oscillospirales genera (Fig. 2, Table S5, S6).

The second type is PEPCK, which has been unpreceded in *cooS* genomic contexts. PEPCK (*pckA*; K01610, COG1866) is a phosphoenolpyruvate carboxykinase that catalyzes the conversion of phosphoenolpyruvate (PEP), CO_2_, and ADP to oxaloacetate (OAA) and ATP, and vice versa. The PEPCK type was found among the genomes of Veillonelales genus *Veillonella* (189/342 genomes). Gene composition varied among PEPCK-type genomic contexts. Of the 189 PEPCK-type genomic contexts of *Veillonella*, 144 included a putative *oxyR*, which encodes a transcriptional regulator of LysR family (K04761, COG0583) and responds to oxidative stress (Sen and Imlay 2021) in the upstream region adjacent to *cooS*. Genes for nitrate reduction (*narGHIJK*) were also observed in 111 *Veillonella* PEPCK-type genomic contexts, of which 78 genomes also included *oxyR* (Fig. 2).

The third is FNOR type that has been predicted to be associated with CO utilization in the previous studies (Geelhoed *et al*. 2016; Inoue *et al*. 2019, 2022). FNOR (COG1251) is often present in the CO utilizers and is involved in energy conservation through CO oxidation associated with the reduction of NAD(P)^+^ (Geelhoed *et al*. 2016). In human gut microbial genomes, FNOR genes were found in 138 *cooS* contexts of bacteria, including Lachnospirales (15/38 genera, 92/379 genomes) and Peptostreptococcales (5/9 genera, 18/41 genomes) (Fig. 2).

The following types have been reported previously; however, their physiological functions in CO metabolism are unknown: ABC transporters (K02049–51, COG0715-COG0600-COG1116) were found in various bacteria, such as Lachnospirales (9/38 genera, 40/379 genomes), Campylobacterales *Campylobacter* (37/40 genomes), and Selenomonadales *Selenomonas* (4/4 genomes) (Fig. 2). “Fe-only hydrogenase” (COG4624) was encoded in Lachnospirales (9/38 genera, 43/379 genomes), Oscillospirales (2/13 genera, 6/411 genomes), and Peptostreptococcus (2/9 genera, 4/56 genomes) (Fig. 2, Table S6). The COG4624 gene was located downstream region adjacent to *cooS* in 24 of the 57 *cooS* genomic contexts with COG4624, including contexts from Lachnospirales *Anaerobutyricum*. In 17 of these, *pflAB* (K04069 and K00656) was found within the genomic context of *cooS* (Fig. 2). In addition to the Fe-hydrogenase-type context, COG4624 was also found in the WLP-type context in 27 genomes.

“MFS transporter,” “functionally uncharacterized dehydrogenase,” and “Cysteine synthase” were relatively minor types, as they occupied fewer than 2% of the CODH-bearing genomes and were observed in specific taxa. The MFS transporter is responsible for importing or exporting a wide range of substrates across the membrane using a substrate concentration gradient. The MFS transporters encoded in this context were annotated as COG2223 (NarK, nitrate transporter) and K08177 (OxlT, oxalate/formate antiporter) in Selenomonadales *Mitsuokella* (16/20 genomes) (Fig. 2). In the functionally uncharacterized dehydrogenase context type, genes for K00004 (butanediol dehydrogenase) and COG1063 (threonine dehydrogenase or related Zn-dependent dehydrogenase) were found in the downstream region adjacent to *cooS* in Clostridiales *Clostridium* (11/15 CODH-bearing genomes), whereas genes for hydrogenase nickel incorporation protein (*hypAB*, K04651, and K04652), which may be involved in maturation of CooS, were present upstream of *cooS*. Cysteine synthase (CysK; K01738, COG0031) was mainly found in Veillonellales *Megasphaera* (11/12 genomes).

### Fdh-lacking WLP is common in the human gut microbiome

As the WLP-type genomic context was the most prevalent among the 1,302 potential CO utilizers in the human gut, we further investigated whether human gut prokaryotes bearing the WLP context were capable of acetyl-CoA synthesis through the WLP. Since WLP could be functional even when *cooS* and *acsB* are not located in the same genomic context (Gencic and Grahame 2020), we here referred to the 667 genomes bearing both *cooS* and *acsB* as CODH/ACS-bearing genomes regardless of whether to retain them as the WLP type context. The presence of other WLP genes such as *cooCF*, *fdh*, *fhs*, *folD*, *metF*, and *acsCDE* was surveyed in all the genomes containing genes for CODH/ACS (667 genomes) (Fig. 3).

**Fig. 3.**
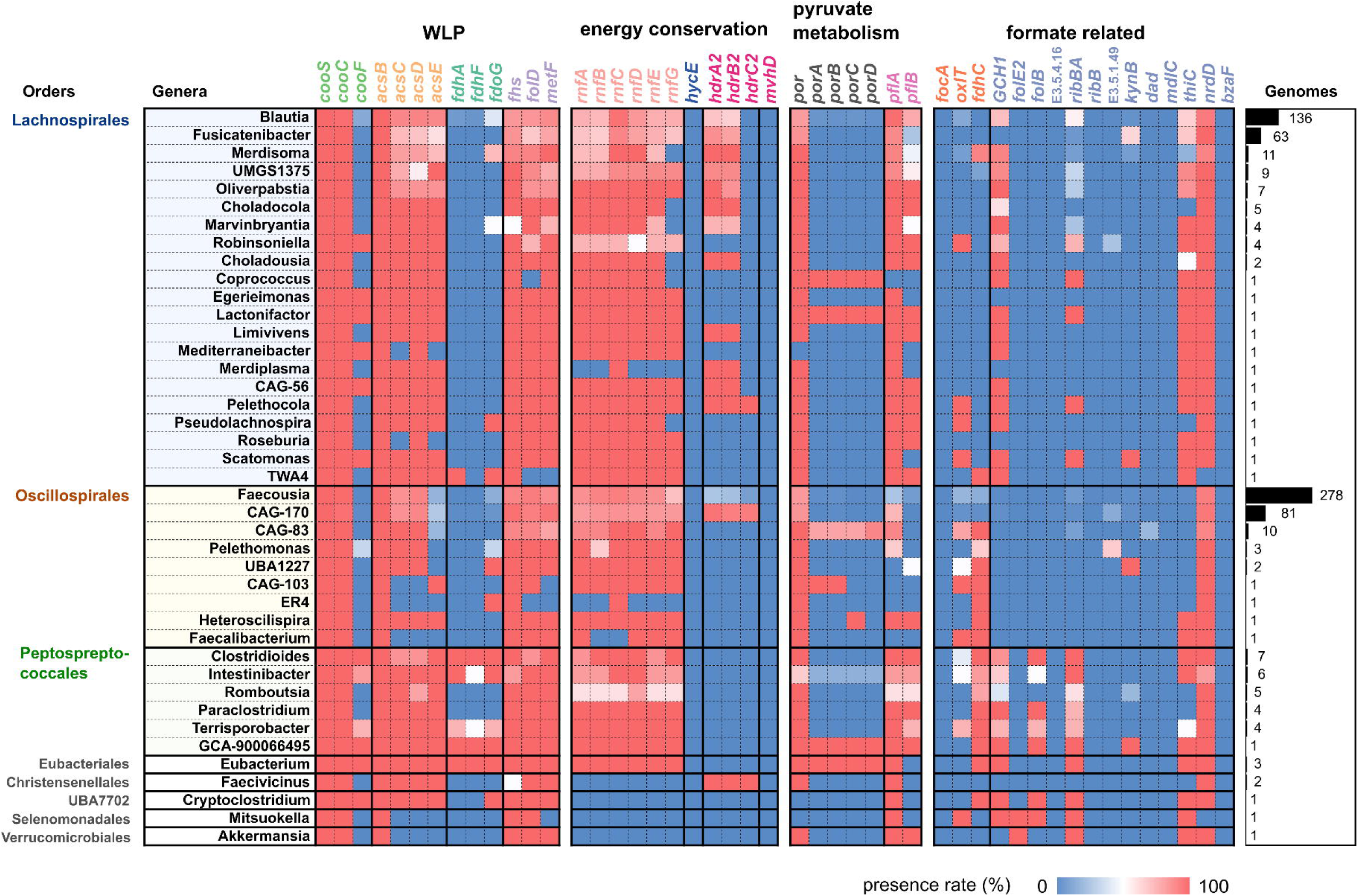
The presence rates of the WLP genes and other related genes in the human gut genomes. Genomes possessing *cooS* and *acsB* were analyzed to reveal the presence of WLP-related and other genes. The presence rates are displayed for each genus on a heatmap. *cooS*, carbon monoxide dehydrogenase catalytic subunit gene; *cooC*, carbon monoxide dehydrogenase maturation gene; *cooF*, ferredoxin-like protein-coding gene; *acsBCDE*, acetyl-CoA synthase gene; *fdhA*/*fdhF*/*fdoG*, formate dehydrogenase catalytic subunit gene; *fhs*, formyltetrahydrofolate synthase gene; *folD*, methylene-tetrahydrofolate dehydrogenase/cyclohydrolase gene; *metF*, methylene tetrahydrofolate reductase gene; *rnf* ferredoxin:NAD^+^ oxidoreductase gene; *hycE*, energy-converting hydrogenase catalytic subunit gene; *hdr*-*mvh,* methylenetetrahydrofolate reductase gene; *por*/*porABCD*, pyruvate: ferredoxin oxidoreductase *pfl*, pyruvate: formate lyase gene. *focA*/*oxlT*/*fdhC*, genes are involved in formate transport. Other genes, genes involved in formate consumption and metabolism. The right bar represents the number of genomes in each genus.

More than 88% of CODH/ACS-bearing genomes retained almost the full set of WLP genes, including *acsCD*, *cooC*, *fhs*, *folD*, and *metF* (Fig. 3, Fig. S2). Interestingly, *acsE*, a bacteria-specific gene for the methyltransferase subunits of ACS, was not detected in 78% of the Oscillospirales genomes (7/9 genera) (Fig. 3), whereas *acsE* was conserved in other CODH/ACS-bearing genomes. Although a nearly full set of WLP genes was detected, we found that the Fdh genes for formate synthesis (*fdhA*, *fdhF*, and *fdoG*) were not detected in 95%, 96%, and 79% of the CODH/ACS-bearing genomes, respectively (Fig. 3), and any other types of the Fdh genes were observed in 79% of the genomes. A previous study reported that two *Blautia* species are capable of CO metabolism in the presence of external formate even though the bacteria lack Fdh gene/activity (Trischler *et al*. 2022). In this study, the presence of Fdh genes were also checked in the 554 CODH/ACS-bearing genomes found in other environments (Inoue *et al*., 2022) (Fig. S2). It revealed that Fdh-lacking WLP is also present in the genomes of other environments. However, compared to human gut microbial genomes with WLP genes, Fdh genes (*fdhA*, *fdhF*, and *fdoG*) were missing in a smaller proportion of bacteria of other environments (28%, 84%, and 17%, respectively) (Fig. S2), and none of the Fdh genes was observed in 14% of the genomes. Thus, the lack of formate synthesis might be a feature highly enriched in the human gut potential CO utilizer. Similarly, *cooF* was not detected in 91% of the CODH/ACS-bearing genomes (accounting for 26/42 genera), whereas *cooF* was absent in 53% of the bacteria isolated from other environments (Fig. 3, Fig. S2).

The above analyses indicate that there may be CO_2_/H_2_-independent sources of formate for human gut CODH/ACS-bearing bacteria. To estimate how formate is obtained, we surveyed the genes involved in alternative formate synthesis or import by CODH/ACS-bearing bacteria (Fig. 3). The Lachnospirales (16/21 genera, 145/239 genomes), a few Oscillospirales genera (2/9 genera), all Peptostreptococcus genera (5/5 genera, 22/26 genomes), and Verrucomicrobiales genera Akkermansia (1 genome) contained *pflB* (Fig. 3). These genera are likely capable of producing formate from pyruvate. Three Oscillospirales genera that lacked both *fdh* and *pfl* possessed a gene for formate transporters, *fdhC* (56/291 genomes) (Fig. 3). Two Lachnospirales (*Merdiplasma* and *Fusicatenibacter*) and two Oscillospirales genera (CAG-170 and *Faecousia*) were not found to possess these genes. These genera possess genes encoding proteins involved in formate-producing/utilizing reactions such as GTP degradation (GCH1, *ribBA*) and thiamine metabolism (*thiC*) (Fig. 3).

To gain deeper insights into the functions specifically encoded in the human gut microbial genomes with WLP, we compared all the genes in the genomes of human gut bacteria and other environmental bacteria that contain WLP genes (Fig. S1, Table S7). We observed that genes involved in carbohydrate and saccharide degradation were more prevalent in the human gut microbiome containing WLP genes than in bacteria of other environments (Fig. S1, Table S7). Therefore, one of the possible explanations is that potential human gut CO utilizers with WLP are capable of utilizing more diverse carbohydrates and/or polysaccharides than the CO utilizers of other environments, producing more pyruvate through glycolysis. For pyruvate metabolism, 53% of the human gut CODH/ACS-bearing genomes possessed *pflB*. PFOR (*por*) is also involved in WLP by supplying reduced ferredoxin from pyruvate oxidation (Ragsdale 2004) and is present in 90% of the human gut CODH/ACS-bearing genomes. These genes were less frequently observed in CODH/ACS-bearing bacteria in other environments: *por* and *pflB* were present in 71% and 19% of the bacteria in other environments, respectively (Fig. S2).

Not only genes for carbohydrate/saccharide degradation (above), but also the genes for Rnf complex (*rnfABCDEG*) were detected in a high proportion of WLP-bearing CO utilizers in the human gut (>73%). Group B FeFe hydrogenase, which is involved in ferredoxin-coupled H_2_ production (Benoit *et al*. 2020), was relatively enriched in human gut WLP-bearing CO utilizers (64%) (Fig. S2). Since some FeFe hydrogenases are known as electron-bifurcating enzyme that provide Fd to Fd-dependent Fdh of CO-utilizing acetogen (Schuchmann and Müller 2012), the co-occurrence of FeFe hydrogenase and Fdh was checked. As a result, group A13, A4, C1, and C3 FeFe hydrogenases were co-occurring with Fdh genes (Fig. S3). Especially, the group A4 FeFe hydrogenase and *fdhA* were statistically co-occurring, with an adjusted odds ratio of 2,376 and a 95% confidence interval ranging from 136 to 41,655 (Fig. S3). Furthermore, *fdoG* was co-occurring with group C3 FeFe hydrogenases, with an odds ratio of 59.4 and a 95% confidence interval ranging from 25.6 to 138 (Fig. S3). However, the most prevalent type of hydrogenase, FeFe groupB hydrogenase gene, showed no association with Fdh genes (Fig. S3).

### CODH transcripts were detected from 97.3% of metatranscriptomic samples

To gain insights into whether active CO-utilizing bacteria are present in the human gut microbiome, metatrascriptome samples from 110 healthy human individuals were analyzed. Following quality control, 7.8 ± 1.2 million reads were obtained per sample on average (Fig. 4a). Filtered reads were then mapped to 1,380 CODHs to identify CODH transcripts in the human gut microbiome. It was found that CODHs were transcribed in 107 out of 110 metatranscriptome samples, with an average of 1,127 reads mapped to the CODH sequences (Fig. 4b). The number of detected CODHs correlated with the total number of reads (*r^2^* = 75.7), with nearly full-length CODH mRNAs (cover length >1500 nt) detected in samples with more than 28 million filtered reads (Fig. 4b). The host bacteria of the CODH transcripts were assigned to the 13 orders from 7 phyla (Fig. 4c). None of the reads from the CODH-bearing genomes of the phyla Campylobacterota, Desulfobacterota, and Proteobacteria were detected in the healthy human metatranscriptome. At the order level, there were Lachnospirales (average 2.9 ×10^5^ RPKM of CODH transcripts across 110 datasets), Oscillospirales (1.7 ×10^5^ RPKM), Peptostreptococcales (2.6 ×10^4^ RPKM), Veillonellales (1.8 ×10^4^ RPKM), Synergistales (1.8 ×10^3^ RPKM), and other 8 orders (55 to 815 RPKM). At the genera level, CODHs from 66 genera were detected (Fig. S4). In Lachnospirales, the genera *Blautia*, *Choladocola*, *Fusicatenibacter* and UMGS1375 accounted for 37.9%, 19%, 12%, and 7.9% of CODH transcripts from Lachnospirales, respectively. In Oscillospirales, the genera *Faecousia* and CAG-170 accounted for 65% and 26% of the transcripts, respectively. In Peptostreptococcales, 58% of CODH transcripts were from the genus *Intestinibacter*. In Veillonellales, 99.7% of CODH transcripts were derived from the genus *Veillonella*. Regarding the function of CODHs, 6 out of 8 types (WLP, PEPCK, FNOR, ABC transporter, Fe-hydrogenase, MFS transporter) were identified in the human gut metatranscriptomes (Fig. 4d). The most abundant type was WLP, accounting for 66% relative abundance on average, while average abundances of other types ranged from 0.02 to 5.0% (Fig. 4d).

**Fig. 4.**
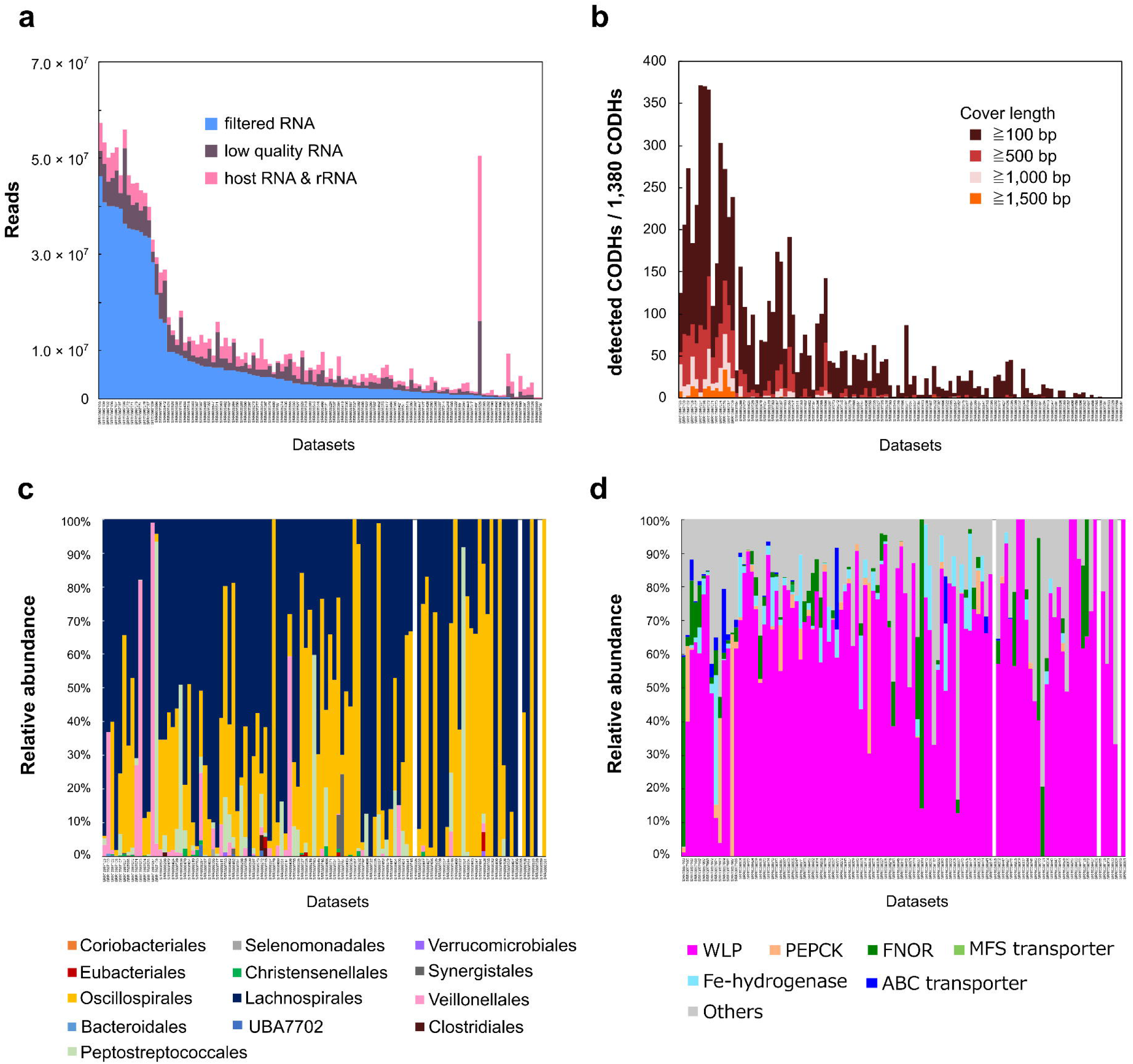
Prevalence and diversity of CODH transcripts in the healthy human gut metatranscriptomic datasets. (a) Composition of sequence reads in the 110 of publicly available metatranscriptomes in the health human stool samples. (b) Number of detected CODHs among 1,380 non-redundant CODH sequences. (c) Relative abundances of host bacteria of the detected CODH sequences in the metatranscriptomic datasets. (d) Relative abundances of CODH types in the metatranscriptomic datasets. In the x-axis of the graphs, datasets with a greater number of filtered RNA are aligned from left to right.

To determine if the entire WLP is transcribed in the human gut microbiome, transcripts of WLP-related genes were investigated. Among the most prevalent WLP-bearing species, genomes with completeness greater than 94% were used as the references for read mapping. As a result, transcripts of all WLP genes (*cooS*, *acsB*, *fhs*, *folD*, *metF*) from six species were detected in the human gut metatranscriptome, clearly indicating that WLP is actively transcribed in the human gut (Fig. 5). Transcripts associated with WLP (*rnf*, *pfor*, *pfl*) were also identified in the samples (Fig. 5). Metatranscriptome reads were mapped to *fdhF* from *B. schinkii* and *B. hydrogenotrophica*, resulting in the detection of *B. hydrogenotrophica fdhF* in 4 out of 110 samples, although the cover length was shorter than 500 nt in all four samples. Since PFOR can supply reduced ferredoxin from pyruvate and PFL can supply formate from pyruvate, our interest lay in the expression balance of PFOR and PFL to predict the metabolism of Fdh-lacking bacteria. Both transcripts for PFOR and PFL were identified (Fig. 5 a-d). PFOR transcripts were generally detected at a higher level than PFL, with an average 19.7-fold difference, while PFL transcripts were observed at higher levels than PFOR in three samples (Fig. 5 a-d).

**Fig. 5.**
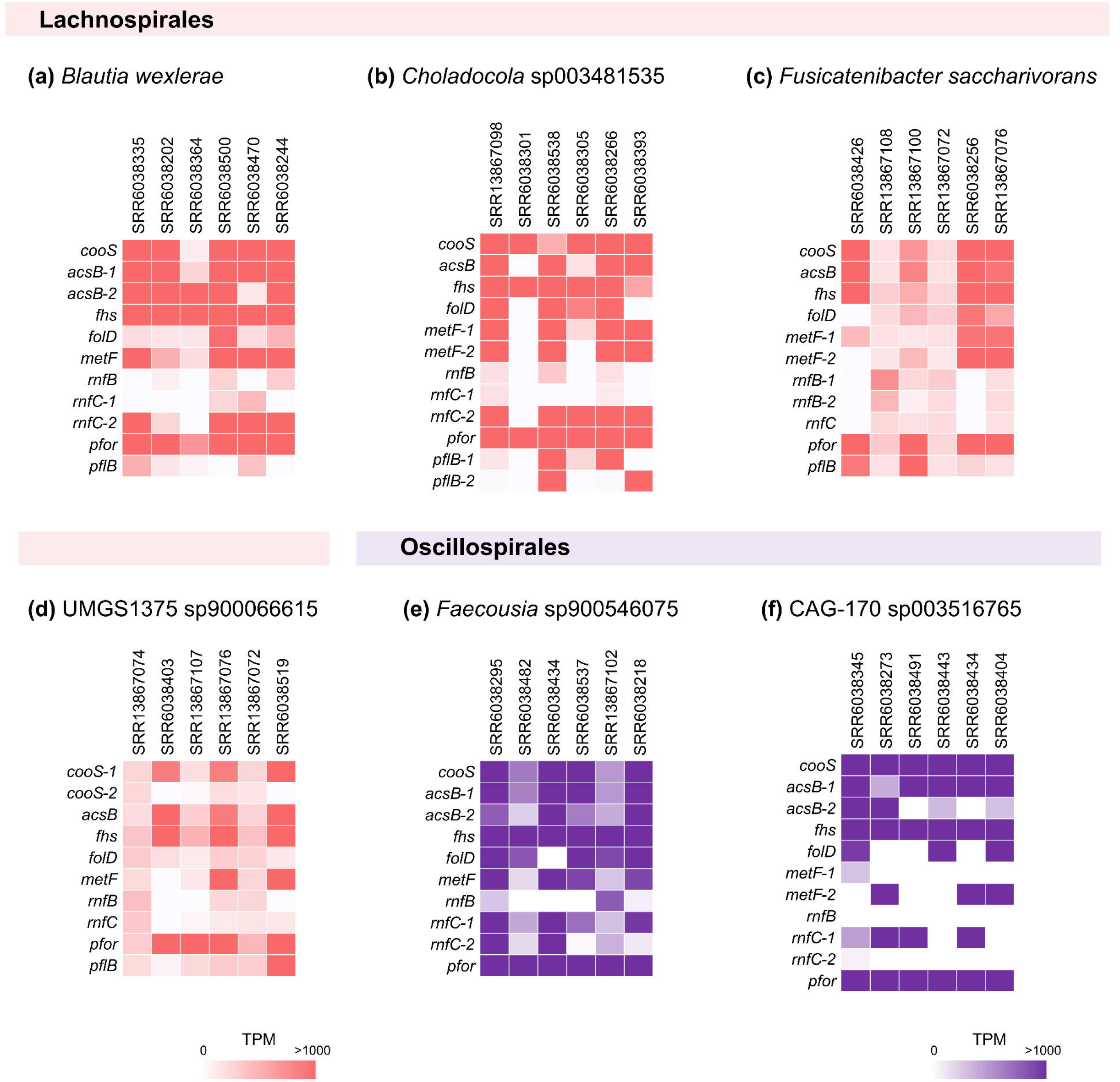
Abundance of WLP-related transcripts in the human gut metatranscriptomes. The heatmaps illustrate the relative abundances of WLP transcripts derived from seven different CODH-bearing genera that were most prevalently detected in the 110 human gut metatranscriptomes. Genomes with more than 94% completeness were selected as references to map the six metatranscriptomic datasets with the highest numbers of CODH reads for the selected genomes. The seven genomes were derived from (a) *Blautia wexlerae*, (b) *Choladocola* sp003481535, (c) *Fusicatenibacter saccharivorans*, (d) UMGS1375 sp900066615, (e) *Faecousia* sp900546075, and (f) CAG-170 sp003516765.

## Discussion

Given the continuous production of CO through heme degradation in the human body (Hopper et al. 2020), it was predicted that a significant number of CO-utilizing prokaryotes exist in the human gastrointestinal tract, contributing to the consumption of the accumulating harmful CO. Previous studies indeed demonstrated the rapid consumption of CO by fresh human fecal samples (Levine *et al*. 1982) and by two *Blautia* strains isolated from human feces in the presence of formate (Trischler *et al*. 2022). However, the mechanism through which taxonomically and phylogenetically diverse human gut prokaryotes become potential CO utilizers in the human gut microbiome remains unclear. In the present study, we analyzed CODH-bearing genomes derived from potential CO utilizers in the human gut microbiome and uncovered various Ni-CODH-bearing genomes belonging to 248 species and 82 genera across 8 phyla (Fig. 1). Furthermore, CODH transcripts derived from 66 genera, 5phyla were identified in 110 of metatranscriptome samples obtained from healthy human with various nationality and age (ranging from 6 months to 75 years old). This suggests that the CO utilizing pathways are active and may constitute an important component of the human intestinal environment. To date, only two human gut prokaryotes have been experimentally verified as CO utilizers (Trischler *et al*. 2022) and 280 *Blautia* genomes and 43 acetogenic species were checked to identify the presence of *cooS* (Trischler *et al*. 2022; Yao *et al*. 2023). Therefore, our findings extensively expand the catalogue of potential human gut CO utilizers.

Our findings are not limited to phylogenetic diversity but include the functional diversity of CO utilizers. The previously identified CO-mediated metabolic pathway in human gut-derived prokaryotes is the WLP (Trischler et al. 2022). However, as their CODH genomic contexts can be divided into eight distinct types presumably involved in distinct physiological roles, the diverse taxa of the human gut microbiome are potentially involved in CO utilization through various pathways.

We observed that WLP is most prevalent in the genomes and metatranscriptome of human gut microbes, such as those of Oscillospirales, Lachnospirales, and Peptostreptococcales. Almost all genes responsible for carbonyl and methyl branches, including a gene for utilizing exogenous CO, were identified in the genomes of the human gut microbiome. However, any types of Fdh genes were not detected in large number of the WLP-bearing human gut microbial genomes (Fig. 3). Previous cultivation experiments have suggested that human gut CO utilizers *Blautia luti* and *B. wexlerae* in the order Lachnospirales lack Fdh activity but possess functional WLP in formate-added medium (Trischler *et al*. 2022). These bacteria therefore utilize extracellular formate. Furthermore, these bacteria possess PFL genes in their genomes which synthesizes formate and acetyl-CoA from pyruvate (Fig. 6). It suggests that glycolysis would become one of the primary sources of carbon and reducing power for the Fdh-lacking WLP in *Blautia luti* and *B. wexlerae* (Fig. 6), meaning that their WLP may be modified to a heterotrophic form that obtains carbon from organic compounds instead of CO_2_. The metabolic pathways of this “heterotrophic WLP” should be tested experimentally. However, the coexistence of PFL and PFOR in *B. luti* and *B. wexlerae* suggests that carbon and electron fluxes may be balanced depending on the available carbohydrates and electron donors in the environment (Fig. 6). Additionally, group B FeFe hydrogenase possessed by these *Blautia* strains may be utilized to dispose of the excess amount of Fd_red_ produced via glycolysis using PFOR as evident from the cultivate experiments of *B. coccoides* demonstrating hydrogen production in the presence of glucose (Liu *et al*. 2015). We further predicted that this surplus Fd_red_ arises due to the saccharide-rich environment of the human gut (Pokusaeva, Fitzgerald and van Sinderen 2011), and part of Fd_red_ is uptaken by Fdh-lacking WLP to produce acetate as well as the hydrogenase.

**Fig. 6.**
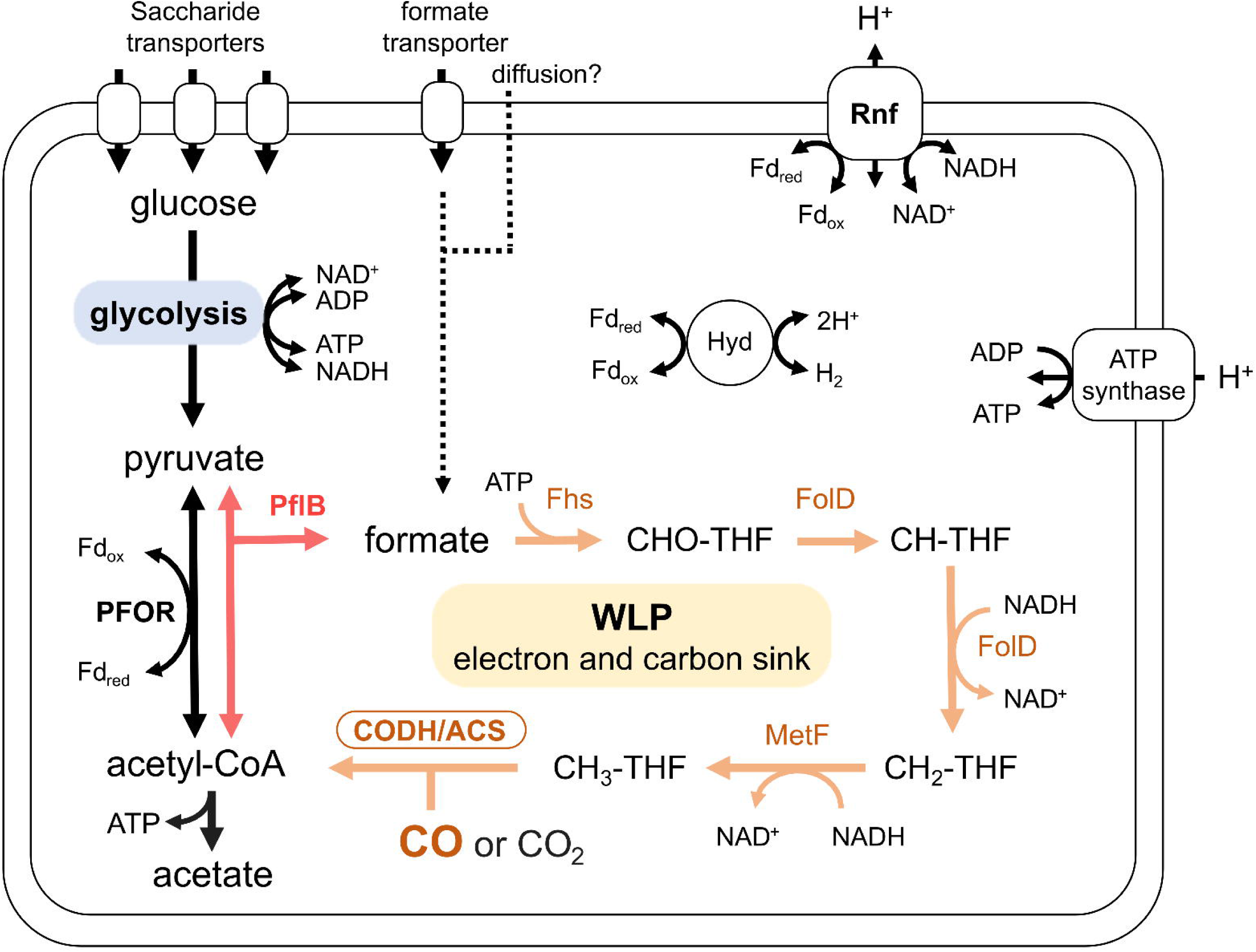
Genome-based metabolic reconstructions of the Fdh-lacking WLP in the human gut. A schematic of Fdh (formate dehydrogenase)-lacking WLP commonly identified in the human gut bacteria, including genera *Blautia* and *Fusicatenibacter* from Bacillota order Lachnospirales. The figure shows an example of metabolic pathways. Pyruvate which is mainly obtained via glycolysis is converted to acetyl-CoA and formate by *pflB* (pyruvate: formate lysase) (red lines). Formate is then progressively converted to methyl moiety in WLP (orange lines) using formyl-tetrahydrofolate synthase (Fhs), methylene-tetrahydrofolate dehydrogenase/cyclohydrolase (FolD), and methylene-tetrahydrofolate reductase (MetF). CO and CH_3_ are then incorporated to acetyl-CoA with CO dehydrogenase/acetyl-CoA synthase (CODH/ACS). The WLP may contribute to toxic CO utilization, carbon acquisition, sink of electrons, and carbons obtained by glycolysis in the human gut prokaryotes.

Although their CO-utilizing ability has not been confirmed, several human gastrointestinal acetogens, including species from Lachnospirales (*B. producta* and *Marvinbryantia formatexigens*) and Clostridiales (*C. bovifaecis* and *Clostridioides difficile*), lack *fdh* but retain WLP genes (Wolin et al. 2003; Yao *et al*. 2020, 2023). This study indicates that the Fdh-lacking WLP are not limited to *Blautia*, *Marvinbryantia*, and *Clostridium* but more prevalently utilized in the human gut CO utilizers represented by the 526 genomes from 32 genera. While a few routes were predicted for formate acquisition, the "heterotrophic WLP" system may represent one favorable pathway, as it is possessed by 53% of WLP-bearing human gut bacteria. Importantly, many of the potential CO utilizers with such “heterotrophic WLP” also retain genes to degrade carbohydrates and polysaccharides to greater extent than the CO utilizers of other environments (Fig. S1, Table S7). The presence of these genes, coupled with the availability of carbohydrates in the human intestinal tract (Pokusaeva, Fitzgerald and van Sinderen 2011), suggests an enhanced provision of substrates for glycolysis, and thereby pyruvate for formate synthesis. This alternative pathway for formate synthesis may be one reason why *fdh* was positively dropped from the genomes of acetogens in the intestinal environment (Yao *et al*., 2023). In addition, the previous findings by Levine *et al*. (1982), which observed higher CO consumption activity of feces samples in the presence of glucose (0.7 mL/h, g feces) than in the absence of glucose (0.2 mL/h, g feces) (Levine *et al*. 1982), might at least partially be caused by the heterotrophic WLP-bearing human gut CO utilizers. Regarding the other routes for formate acquisition, 141 genomes possessed Fdh genes. These Fdh genes were found to co-occur with putative electron-bifurcating group A13, A4, C1, and C3 hydrogenases (Fig. S3), a condition necessary for utilizing CO to prevent toxicity to the hydrogenase module of H_2_-dependent CO_2_ reductase (HDCR) (Dietrich and Müller 2023). There were also bacteria lacking both Fdh and PFL. These bacteria may import formate from extracellular fractions by using the formate transporters or by cross-feeding (Gomez *et al*. 2019).

A novel CODH type, which has not been previously reported, was identified in 190 genomes of the genus *Veillonella* (Fig. 2). *Veillonella* mainly harbored clade B CODH, a phylogenetically separate variant from the predominant WLP-associated clade E CODH (Fig. 1). The *cooS* genome of *Veillonella* contained genes encoding PEPCK and NarKGHJI (Fig. 2), which may be involved in CO metabolism. For instance, a biochemical link between CO oxidation and nitrate reduction may be established as observed in *Deferribacter autotrophicus* (Slobodkin *et al*. 2019). However, as the reconstructed pathways described above are based only on metagenome-assembled genomes and metatranscriptomes, further experimental verifications are necessary.

This study revealed the phylogenetic diversity of CO utilizers and their divergent CO-utilizing pathways in the human gut microbiome. Because trace amounts of CO are constantly produced in the human body, CO utilizers may constantly consume CO in the human gut microbiota, avoiding the accumulation of high concentrations of toxic CO in the gut. Therefore, shedding light on CO utilizers would help elucidate the microbial mechanisms underpinning intestinal homeostasis in humans.

## Acknowledgments

Computation time was provided by the Super Computer System, Institute for Chemical Research, Kyoto University.

## Author contributions

**Yuka Katayama**: Conceptualization, Methodology, Software, Validation, Formal analysis, Writing - Original Draft, Writing - Review & Editing, Visualization. **Ryoma Kamikawa**: Conceptualization, Writing - Original Draft, Writing - Review & Editing, Supervision. **Takashi Yoshida**: Conceptualization, Writing - Review & Editing, Supervision, Funding acquisition.

## Figure legends

**Fig. S1.**
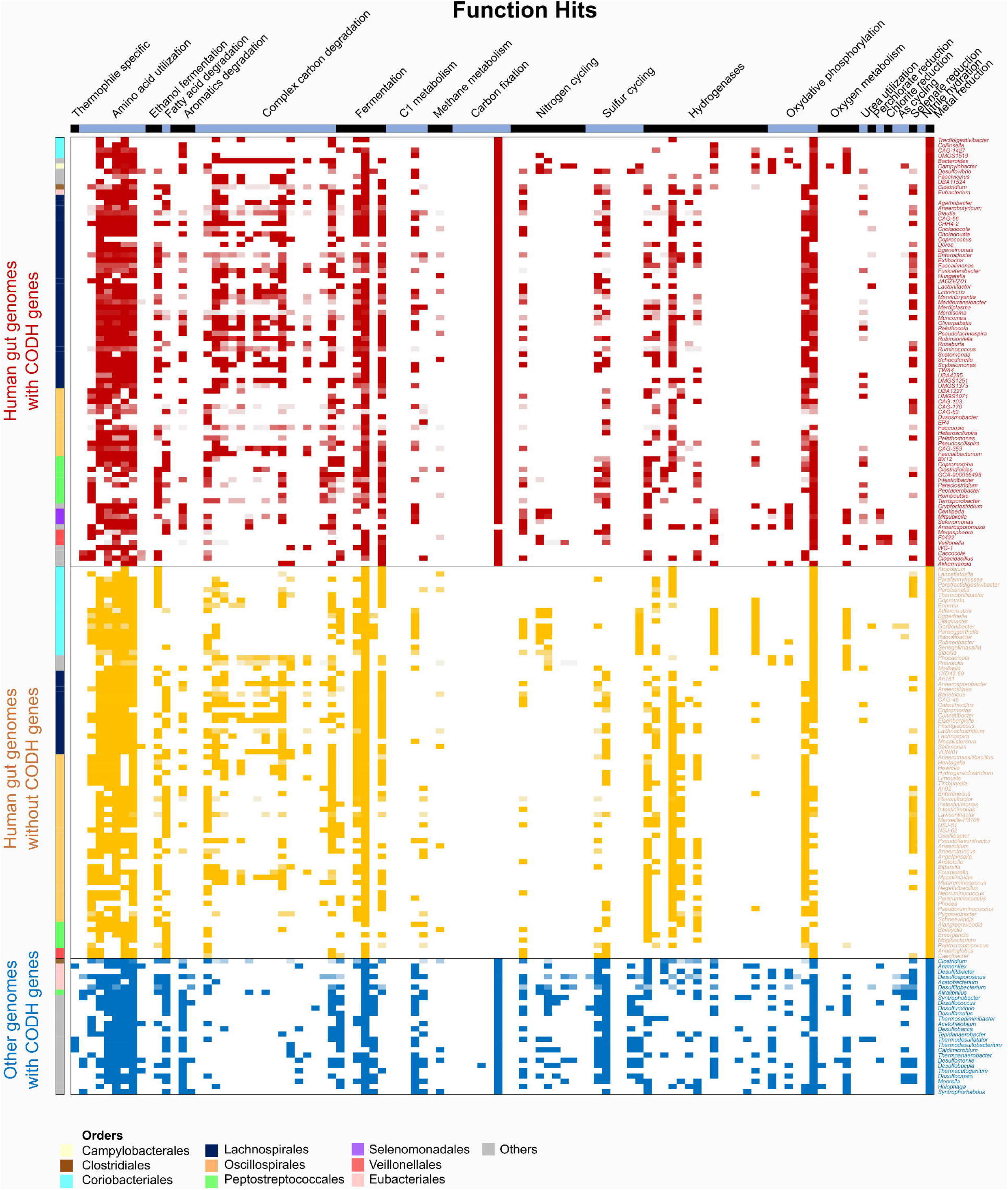
Metabolic function hits detected in the three groups of genomes. The metabolic functions of the three genome groups were analyzed using METABOLIC v4.0. The y-axis represents the genomes grouped by phylogeny (genera and orders) and origin. The first group consists of CODH-bearing genomes detected in the HumGut database (red). The second group comprises non-CODH-bearing genomes, which belong to the same family as the CODH-bearing genomes in the HumGut database (yellow). The third group comprised CO utilizers found in environments other than that of the intestines (blue). The x-axis represents the functions identified in each genome group. The functions were characterized by the presence of genes in METABOLIC. The following list represents the functions and the detected genes associated with each function, which are ordered from left to right in the columns: thermophilic-specific (reverse gyrase), amino acid utilization (4-aminobutyrate aminotransferase and related aminotransferases, aminotransferase class I and II, phosphoserine aminotransferase, ornithine/acetylornithine aminotransferase, branched-chain amino acid aminotransferase/4-amino-4-deoxychorismate lyase, aspartate/tyrosine/aromatic aminotransferase, histidinol-phosphate/aromatic aminotransferase, serine-pyruvate aminotransferase/archaeal aspartate aminotransferase), ethanol fermentation (aldehyde dehydrogenase, alcohol dehydrogenase), aromatics degradation (*catA*, *ubiX*||*bsdC*, *bcrABCD*), complex carbon degradation (cellobiosidase, cellulase, beta-glucosidase, arabinosidase, beta-glucuronidase, alpha-L-rhamnosidase, mannan endo-1,4-beta-mannosidase, alpha-D-xyloside xylohydrolase, beta-xylosidase, beta-mannosidase, beta-galactosidase, alpha-amylase, glucoamylase, pullulanase, isoamylase, chitinase, hexosaminidase), fermentation (*porA*, *adh*, *ldh*, *acdA*||*ack*||*pta*, *acs*, *pflD*), C1 metabolism (*mxaF* or *mdh*, *mauAB*, *fdhA*||*fghA*||*frmA*||mycoS_dep_FDH||*fae*, *fdoG*||*fdwB*||*fdoH*||*fdhAB*, *coxS*||*coxM*||*coxL*), methane metabolism (*pmoABC*, *mmoBD*, *mcrABC*), carbon fixation (Form I, Form II, *mcr*||K14469, K14466||K18861, K18861||4hbl, *cdhD*||*cdhE*||*cooS*, *aclAB*), nitrogen cycling (*amoABC*, *anfDKG*||*nifDK*||*vnfDKG*||*nifH*, *nxrAB*, *napAB*||*narGH*, *nrfADH*||*nirBD*, *nirKS*||*octR*, *norBC*, *nosDZ*, *hzoA*||*hzsA*), sulfur cycling (*fccB*||*sqr*, *dsrABD*||*asrABC*, *sdo*||*sor*, *sreABC*||*sor*, *soxBCY*, *aprA*||*sat*, *phsA*), hydrogenase (FeFe-group-a13, FeFe-group-a2, FeFe-group-a4, FeFe-group-b, FeFe-group-c1, FeFe-group-c2, FeFe-group-c3, Fe hydrogenase, Nife-group-1, NiFe-group-2ade, NiFe-group-2bc, NiFe-group-3abd, NiFe-group-3c, NiFe-group-4a-g, NiFe-group-4hi), oxidative phosphorylation (*nuoABC*, *ndhABC*, *sdhCD*, *petAB*||*fbcH*, *atpAB* (V/A-type), *atpAD* (F-type)), oxygen metabolism (*coxAB*, *ccoNOP*, *cyoABCD*, *cydAB*, *qoxAB*), urea utilization (*ureABC*), halogenated compound utilization (E3.8.1.2||*pcpC*||*cprA*||*pceA*), perchlorate reduction (*pcrAB*), chlorite reduction (*cld*), arsenic cycling (*arxA*||*aioA*, *arrA*), selenate reduction (*ygfMK*||*xdhD*), nitrile hydration (*nthAB*), and metal reduction (iron reduction series genes). Only *acdA*||*ack*||*pta* for fermentation and *lacZ* for complex carbon degradation are highlighted.

**Fig. S2.**
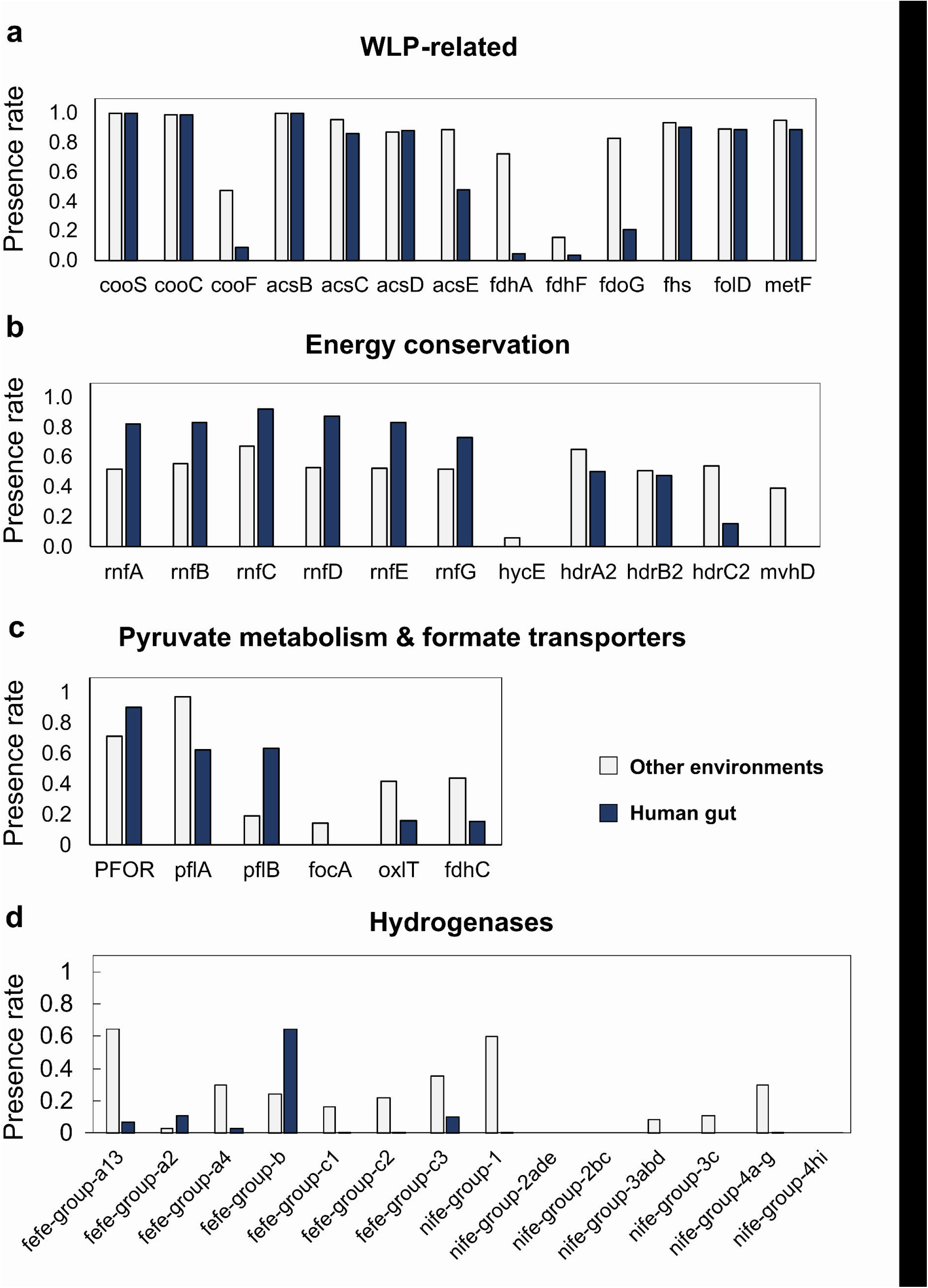
Comparisons of the gene presence rates between human gut and other environmental WLP-containing bacteria. The presence of WLP-related genes was compared between the genomes 667 of human gut bacteria and 554 bacteria from other environments, all of which possess CODH/ACS. The genes were classified into four categories: (a) WLP-related genes, (b) genes involved in energy conservation, (c) genes involved in pyruvate metabolism, and (d) hydrogenase genes. The presence of these genes was determined by analyzing KO annotations using eggNOG-mapper v2.2.5. The presence of hydrogenase genes was assessed using METABOLIC v4.0. *cooS*, carbon monoxide dehydrogenase catalytic subunit gene; *cooC*, carbon monoxide dehydrogenase maturation gene; *cooF*, ferredoxin-like protein-coding gene; *acsBCDE*, acetyl-CoA synthase gene; *fdhA*/*fdhF*/*fdoG*, formate dehydrogenase catalytic subunit gene; *fhs*, formyltetrahydrofolate synthase gene; *folD*, methylene-tetrahydrofolate dehydrogenase/cyclohydrolase gene; *metF*, methylene tetrahydrofolate reductase gene; *rnf*, ferredoxin:NAD^+^ oxidoreductase gene; *hycE*, energy-converting hydrogenase catalytic subunit gene; *hdr*-*mvh,* methylenetetrahydrofolate reductase gene; *porABCD*, pyruvate ferredoxin oxidoreductase gene; *pfl*, pyruvate: formate lyase gene; *focA*/*oxlT*/*fdhC*, genes involved in formate transport. Other genes, genes involved in formate consumption and metabolism.

**Fig. S3.**
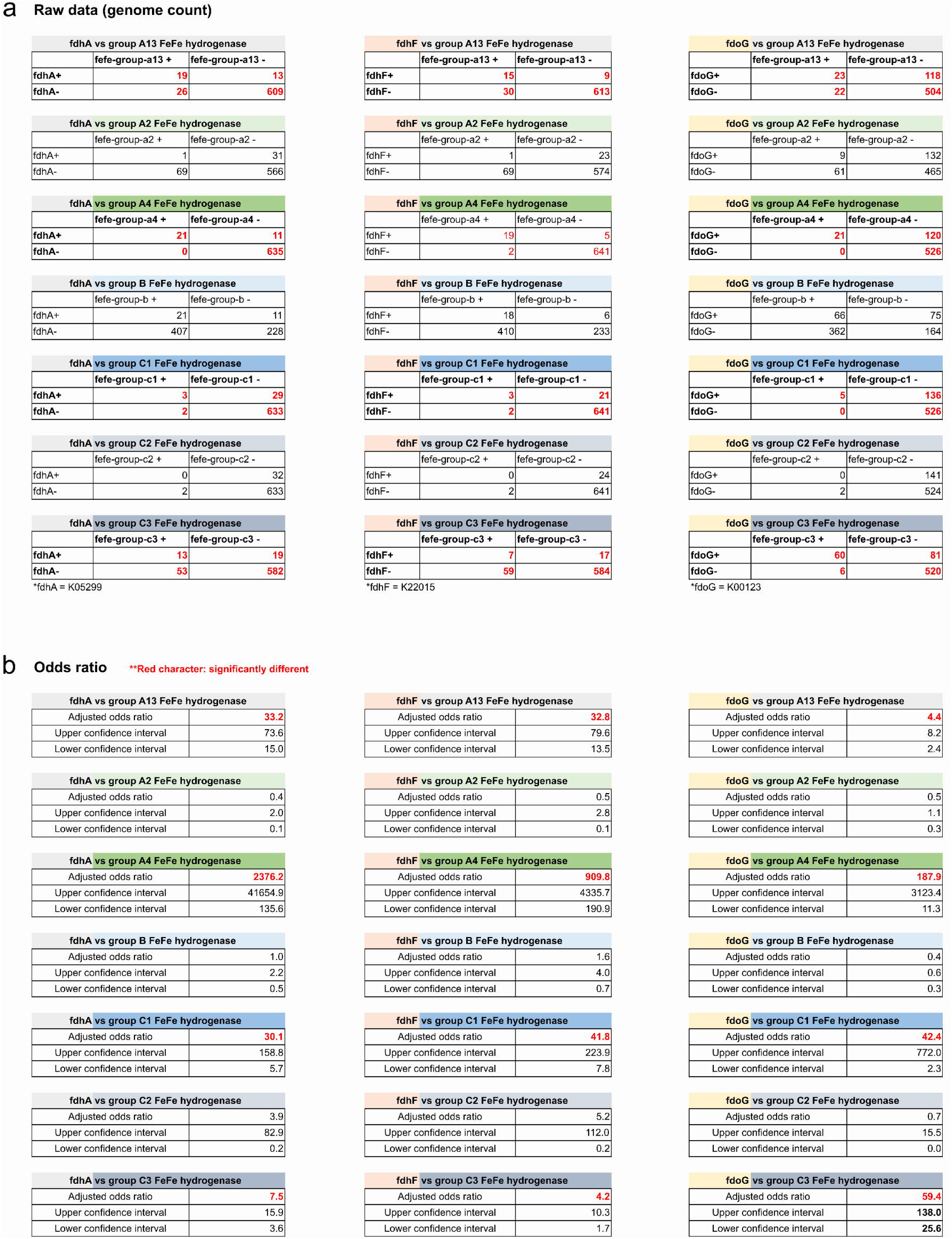
The co-occurrence of genes for Fdh and FeFe hydrogenase in the genomes of CODH/ACS-bearing bacteria in the human gut. (a) The count of genomes containing Fdh genes and FeFe hydrogenase genes, as well as those lacking either gene. These counts were utilized to calculate odds ratio. (b) The odds ratio for the co-presence of Fdh and FeFe hydrogenase genes. The red characters indicate there were significant differences.

**Fig. S4.**
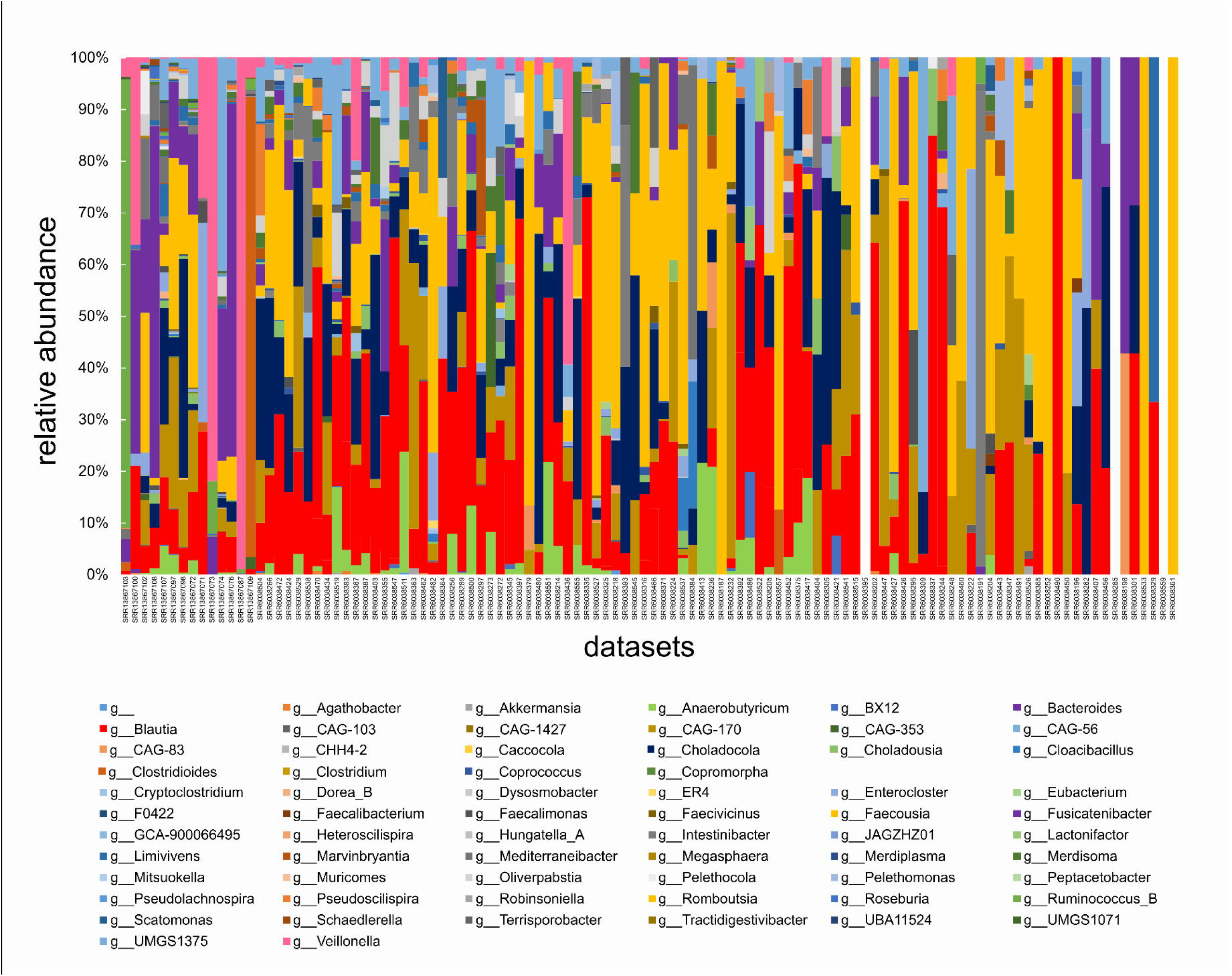
Genera level diversity in the healthy human gut metatranscriptomic datasets. The genera level classification of the hosts of detected CODH transcripts are displayed. In the x-axis of the graph, datasets with a greater number of filtered RNA are aligned from left to right.

